# Water-saving GC-MC model captures temporally differential enzymatic and transporter activities during C3-CAM transition

**DOI:** 10.1101/2024.09.02.610818

**Authors:** Devlina Sarkar, Sudip Kundu

## Abstract

CAM (Crassulacean Acid Metabolism) plants reduce the water loss through transpiration in arid environments (Wickell et al., 2021), using an alternative pathway of carbon assimilation. To ensure food security, engineering CAM into C_3_ plants can be achieved by inverting the stomatal rhythm and the timing of major CO_2_ uptake from day to night. Identification of the metabolic enzymes and intra-cellular transporters, present in both C_3_ and CAM but having different differential temporal activities throughout the diel cycle (Yang et al., 2015; Heyduk et al., 2019) and quantitative estimations of the flux distributions along the biochemical trajectory of C_3_-to-CAM transition may help us to achieve the goal. Here, we simulate a constraint-based combined metabolic model of guard cell (GC) and mesophyll cell (MC), linking temporal fluctuations of temperature (T) and relative humidity (RH) throughout the diurnal cycle with osmolyte accumulation dependent stomatal opening, CO_2_ uptake and transpirational water loss. Starting with C_3_ metabolism, gradual increase in water-use efficiency (WUE) captures several known and new differential activities of metabolic enzymes, transporters, sugar-malate cycle etc. in GC and MC during C_3_-to-CAM transition.

---

Previous computational studies compared the MC metabolisms of C_3_ and CAM plants (Cheung et al., 2014), explored the potential metabolic C_3_-to-CAM transition of MC by declining environmental CO_2_ uptake at daytime (Tay et al., 2021), analysed metabolic changes in MC during C_3_-to-CAM transition by reducing transpirational water loss while compromising phloem output (Töpfer et al., 2020), and studied the C_3_ GC metabolism (Tan and Cheung, 2020). However, metabolisms of GC and MC are interconnected. The stomatal pores, regulated by the osmolytes accumulation in GC, control the transpirational water loss and the uptake of CO_2_. While this CO_2_ is the source of carbon for both GC-MC metabolisms, MC supplies sucrose, one of the osmolytes, balancing the osmotic pressure (OP) in GC. Thus, investigating the changes in this coupled metabolism as WUE increases, is particularly important. Our previously reported six-phase combined metabolic model of GC and MC (Sarkar and Kundu, 2024) has been modified to link the transpirational water loss by the system with the CO_2_ demand, T and RH using a previously used gas diffusion model (Töpfer et al., 2020). Furthermore, to include the occurrence of many (270 to 400) MCs per GC in a leaf (Reckmann et al., 2020), we consider an intermediate value. Firstly, when we maximize the WUE of the model with C_3_ constraints, results connate the switching of the C_3_ metabolism to CAM. Then, we gradually decrease the total water loss throughout the diel cycle and simulate the model using a two-step iteration method to reach to CAM from C_3_ (**Figure 1a**). The increased WUE leads to gradual but nonlinear decrease in daytime CO_2_ uptake, increase in daytime starch and night time malate storage in both GC and MC (**Figure 1b**). Results capture the temporal differences in accumulation of osmolytes like K^+^ (**Figure 1c**) and distinct metabolic behaviours of cells even between different phases of day and night. During the transition, phases 5 and 6 show increased activities of cytosolic PEPC, MDH, and glycolytic enzymes, while phases 1 and 2 exhibit prominent activities of PEPCK, ME, and enzymes related to gluconeogenesis and starch production in both the cells (**Figure 1d-i**). The observed higher starch storage in GC compared to MC at daytime, is also observed in a previous study (Castano, 2020). The enzymatic reactions and transporters having flux values highly correlated with increasing WUE involve different metabolic pathways such as Calvin cycle, gluconeogenesis, glycolysis, TCA cycle, pentose-phosphate pathway (PPP), nitrogen assimilation pathway, mitochondrial and plastidial electron transport chain (ETC), carboxylation-decarboxylation, plastidial-cytosolic and mitochondrial-cytosolic shuttles etc.; and interestingly, the similar patterns for some of them are observed in experimental studies on ice plant by introducing salinity or draught stresses (Kong et al., 2020; Guan et al., 2021). The up- and down-regulated activities of enzymes of above-mentioned pathways throughout the diel cycle are shown in **Figure 2**. Santelia and Lawson, 2016 reported that malate can contribute between 50-90% as a counterion for K^+^ and the result represented here is for maximum value. However, almost similar results are observed within 27-100% and below this, PEPc exhibits higher activity in MC than in GC during daytime in C_3_ plants, which is unlikely to happen (Daloso et al., 2017). The simulation using T and RH of a very hot and dry Indian region, Jaipur indicates the transpirtional water loss varies during C_3_-to-CAM transition, but the pattern remains almost same (**Supplementary File**). In summary, our time-resolved combined model of GC and MC allows us to infer that besides MC, GC also has its own pattern of shifting the metabolism from C_3_ to CAM and same set of enzymes have different temporal activities in the two cells throughout the diel cycle. Although, many physiological and anatomical differences in C_3_ and CAM, like increased succulence in leaf and stem tissues, signaling pathways regulating stomatal rhythm etc., (Griffiths H, 1989; Nelson et al., 2005) are not included in this study, the identification of temporal differential flux pattern of key metabolic pathways and their quantitative estimations add an important step towards engineering CAM into C_3_ plants.

**Figure 1.**
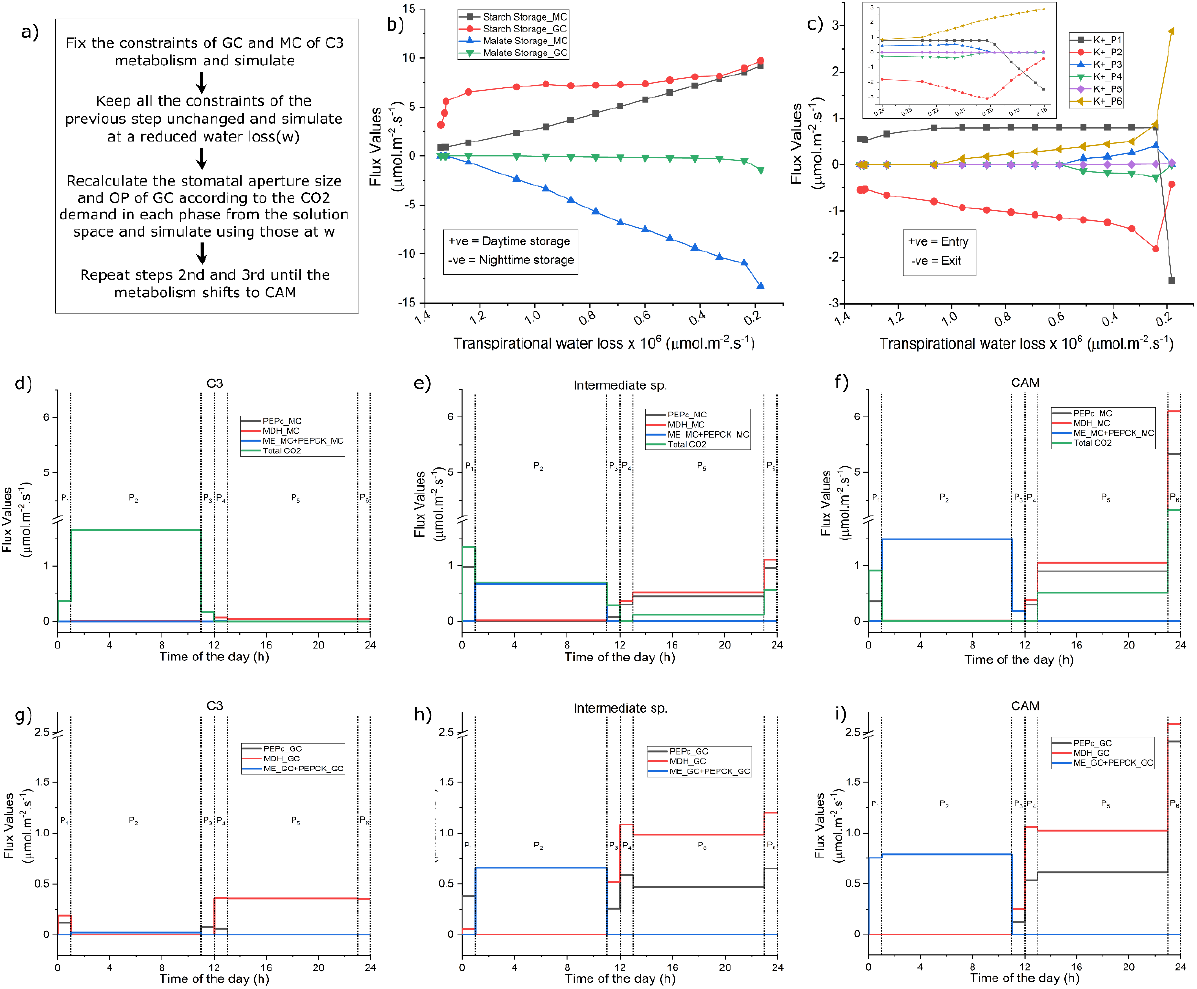
a) The flowchart of the method to simulate C_3_-CAM transition by reducing water loss, b) storage of starch and malate in GC and MC, c) variations in K^+^ uptake and release in GC. Temporal variations of total CO_2_ uptake, activities of PEPc, cytosolic MDH and sum of ME and PEPCK in MC of d) C_3_, e) an intermediate species and f) CAM, activities of these enzymes in GC of g) C_3_, h) an intermediate species and i) CAM are shown. For C_3_ and CAM, the total water loss for 1 GC and 300 MCs are 13,42,711 and 1,80,280 µmol.m^-2^.s^-1^ respectively. An intermediate point of the transition, having a total water loss of 7,80,159 µmol.m^-2^.s^-1^ represents an intermediate species. In b and c, transpirational water loss is presented for 300 MCs and 1 GC. Dotted lines represent the separation of the phases (For details and abbreviations see **Supplementary File**).

**Figure 2.**
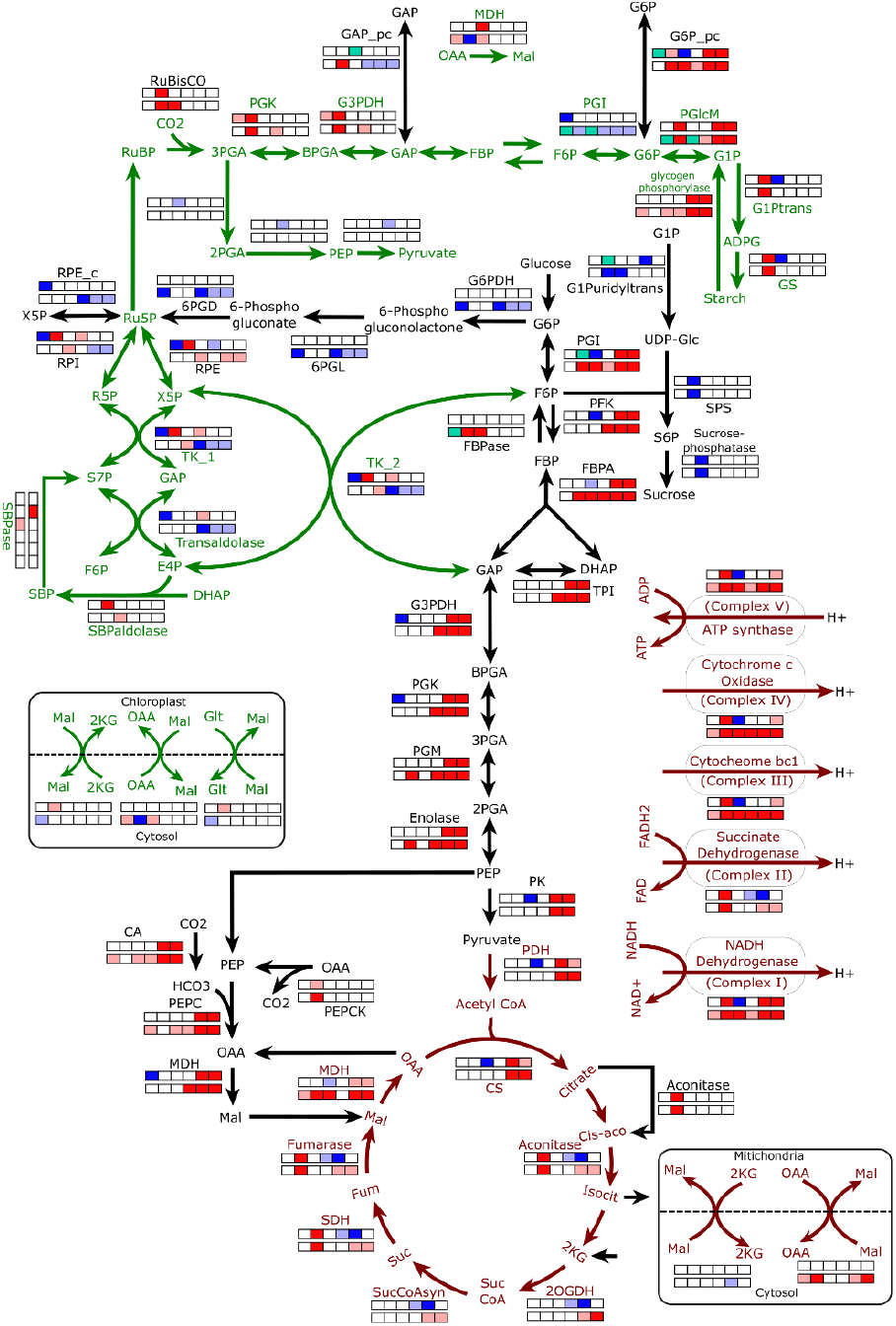
The diagram represents pathways of central carbon metabolism. For each twelve-square box, the upper and lower rows (each row contains six boxes) represent differential activities of enzymatic reactions/transporters in six phases in GC and MC respectively. Red (up-regulation) and dark-blue (down-regulation) represent enzymatic activities having high correlation value (>0.9) with increasing WUE, whereas pink (up-regulation) and light-blue (down-regulation) represent those having correlation between 0.8 and 0.9. The sea-green colour represents that the reaction’s direction changes during C_3_-to-CAM transition in that phase. Green, black and brown arrows represent reactions occurring in chloroplast, cytosol and mitochondria respectively (For details and abbreviations see **Supplementary File**).

## Acknowledgements

DS thanks Department of Biotechnology, Government of India for her fellowship.

## Author contributions

DS and SK designed the study, DS performed the simulations, both interpreted the results and co-wrote the paper.

## Data availability

The model files (in sbml and excel format) are provided in **Supplementary Data S1** and the codes will be provided upon request.

## Supporting information

**Supplementary File**.

**Supplementary Data S1**. The model files in sbml and excel format.

**Supplementary Data S2**. The excel file containing the solution space of each point in the C_3_-to-CAM continuum.

**Supplementary Figure S1**. Plot of the diffusion coefficient of CO_2_ 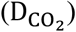 vs temperature (T).

**Supplementary Figure S2**. Schematic representation of the model.

**Supplementary Table S1**. Constraints applied to GC and MC throughout the C_3_-CAM transition.

