## Supplementary File for "Water-saving GC-MC model captures temporally differential enzymatic and transporter activities during C3-CAM transition"

### Abbreviations used in Figure 1 and 2

2KG, 2-Ketogluterate; 2OGDH, 2-oxogluterate dehydrogenase; 2PGA, 2-Phosphoglycerate; 3PGA, 3-Phosphoglycerate; 6PGD, 6-phosphogluconate dehydrogenase; 6PGL, 6-phosphogluconolactonase; BPGA, 1,3-Bisphosphoglycerate; CA, Carbonic anhydrase; CS, Citrate synthase; DHAP, Dihydroxyacetone phosphate; E4P, Erythrose-4-phosphate; FBP, Fructose-1,6-bisphosphate; FBPA, Fructose-1,6-bisphosphate aldolase; FBPase, Fructose-1,6-bisphosphatase; Fum, Fumarate; G1P, Glucose-1-phosphate; G6P, Glucose-6-phosphate; Glc, Glucose; F6P, Fructose-6-phosphate; G6PDH, Glucose-6-phosphate dehydrogenase; G3PDH, Glyceraldehyde-3-Phosphate Dehydrogenase; G1Ptrans, Glucose-1-phosphate adenylyltransferase; G1Puridyltrans, Glucose-1-phosphate uridyltransferase; GAP, Glyceraldehyde-3-phosphate; Glt, Glutamate; GS, Glycogen synthase; Mal, Malate; MDH, Malate dehydrogenase; ME, Malic enzyme; OAA, Oxaloacetate; PDH, Pyruvate dehydrogenase; PEP, Phosphoenolpyruvate; PEPC, Phosphoenolpyruvate carboxylase; PEPCK, Phosphoenolpyruvate carboxykinase; PFK, Phosphofructokinase; PGI, Phosphoglucoisomerase; PGlcM, Phosphoglucomutase; PGK, Phosphoglycerate kinase; PGM, Phosphoglycerate mutase; PK, Pyruvate kinase; RuBP, Ribulose-1,5-bisphosphate; RuBisCO, Ribulose-1,5-bisphosphate carboxylase/oxygenase; RPE, Ribulose-5-phosphate 3-epimerase; RPI, Ribose-5-phosphate isomerase; Ru5P, Ribulose 5-phosphate; R5P, Ribose 5-phosphate; S7P, Sedoheptulose 7-phosphate; SBP, Sedoheptulose-1,7-Bisphosphate; SBPase, Sedoheptulose-1,7-Bisphosphatase; SBPaldolase, Sedoheptulose-1,7-Bisphosphate aldolase; S6P, Sucrose-6-phosphate; SDH, Succinate dehydrogenase; SPS, Sucrose-phosphate synthase; Suc, Succinate; SucCoA, Succinyl-CoA; SucCoAsyn, succinyl-CoA synthetase; TA, Transaldolase; TK, Transketolase; TPI, Triose-phosphate isomerase; X5P, Xylulose-5-Phosphate.

### Details related to Figure 1a

**Figure 1a** shows the 2-step iteration method using which we simulate the $C_3$ to CAM transition. In this study, we have used our previously constructed combined metabolic model of GC and MC (Sarkar and Kundu, 2024) with some necessary modifications to include transpirational water loss, couple this water loss with environmental $CO_2$ exchange, correction of osmotic pressure at different phases of the diel cycle etc. The six phases are named as phase 1 to phase 6 to represent dawn, mid-day, afternoon, dusk, midnight and end of night respectively. The total time of 24 hours of a complete diel cycle is distributed in these six phases in a ratio of 1:10:1:1:10:1. The diffusion flux of $CO_2$ through a stomatal pore can be calculated using the Fick's law of diffusion previously used in Töpfer et al., 2020.

$$J_{CO_2} = D_{CO_2} \frac{n\pi R^2}{l + R_{mean}} \left(c_{CO_2,in} - c_{CO_2,out}\right) \tag{1}$$

Where, $D_{CO_2}$ is the diffusion coefficient of $CO_2$, $c_{CO_2,in}$ and $c_{CO_2,out}$ are inside and outside concentrations of $CO_2$, n is the number of stomata per unit area of the leaf , $\pi R^2$ is the average area per stomatal pore, $l$ is the depth of stomatal pore and $R_{mean}$ is the mean radius which is added to the pore depth as a correction factor (Nobel, 2020). $(c_{CO_2,in} - c_{CO_2,out})$ is considered to have constant value of 0.0055 mol.m$^{-3}$ air as used in Töpfer et al., 2020. The values of $D_{CO_2}$ for distinct temperatures are given in Appendix I in Nobel, 2020. Plotting those points, we get a straight-line graph showed in **Supplementary Figure S1**. Using linear regression method, we get the c (intercept) and m (slope) of the straight line. Now, $D_{CO_2}$ is calculated for the respective temperature of each phase. The aperture size of stomata (R) at day and night are 7 and 0.6 µm (Reckmann et al., 1990) respectively and depth of the stomata is twice the radius of the pore, assuming the pore to be circular (Fanourakis et al., 2015). n is considered to be 700 mm$^{-2}$, within the reported values ranging from 45 to 720 mm$^{-2}$ (Hetherington and Woodward, 2003).

Here, we have linked the metabolisms of GC and MC in such a way that when osmolytes accumulate in GC, stomata open; and depending on the aperture size and diffusion coefficients at different temperatures in different phases of day and night, a maximum amount of $CO_2$ (Calculated using equation 1) can enter the system. We have considered the ratio of GC and MC to be 1/300, an intermediate value in the range of 1/270 to 1/400 (Reckmann et al., 1990). Thus, the entered $CO_2$ is available to 300 MCs and 1 GC. As the phases have different lengths, the amount of $CO_2$ calculated for each phase is further adjusted by multiplying it by the duration of that phase. This $CO_2$ is taken by both GC and MC for their metabolism and on the other hand, MC supplies sucrose, a crucial osmolyte for balancing the osmotic pressure, to GC. $K^+$ can enter and exit GC in any phase of day and night. However, as it is experimentally reported that in GC, $K^+$ accumulates at the early morning and the sucrose accumulation occurs in the later phase of daytime (Lawson, 2009), transfer of $K^+$ from phase 2 to phase 3 is blocked and sucrose is allowed to transfer from MC to GC in phase 2 at daytime. Whereas, at nighttime sucrose is transferred in phase 5. Assuming the photosynthetic capacity of GC to be 20%, the amount of sucrose transferred to GC is 80% of the OP (as in Tan and Cheung, 2020). **Supplementary Figure S2** depicts the schematic representation of the model and the model files can be found in the **Supplementary Data S1**.

Now, the solute content of GC for an open stoma, the initial opening rate of stomata are reported to be 2.02 posmol.cell$^{-1}$ and 3.4 µm.h$^{-1}$ (Reckmann et al., 1990) and the average length and width of guard cells are reported to be 20µm and 9µm (Melaragno et al., 1993). These give us the rate of solute accumulation at daytime to be-

$$\frac{\text{Solute content} * \text{stomatal opening rate}}{\text{aperture size}} = \frac{0.9811\ \text{posmol}}{\text{cell} * \text{hour (h)}}$$

Considering the shape of the GC to be elliptical, we can calculate the rate of solute accumulation per m$^{-2}$ per s.

$$\frac{0.9811 \text{ posmol}}{\text{cell} * \text{h}} = \frac{0.9811 * 10^{-6} \text{ µosmol}}{\pi * \left(\frac{20}{2}\right) * \left(\frac{9}{2}\right) * \ 10^{-12} \text{ m}^2 * 3600\text{s}} = 1.928 \text{ µosmol. m}^{-2}\text{. s}^{-1}$$

Assuming the linear relationship between the OP and aperture size (Reckmann et al., 1990), the OP at night is calculated to be 0.165 µosmol.m$^{-2}$.s$^{-1}$.

$$J_{H_2O} = k\left(c(T)^*_{in,H_2O} - c(T)^*_{out,H_2O}.RH_{out}\right).J_{CO_2} \tag{2}$$

$$\text{where, } k = \frac{D^0_{H_2O}}{D^0_{CO_2}}.\frac{1}{c_{in,CO_2} - c_{out,CO_2}}$$

Equation 2 gives us the relationship between the transpiratinal water loss and $CO_2$ demand of the system depending on the temperature and relative humidity in each phase throughout the diel cycle. T and RH are the temperature and relative humidity at any phase, generated by the skewed sinus generator using the parameters from Töpfer et al., 2020 and maximum and minimum values of T and RH of a diel cycle. $c(T)^*_{H_2O}$ is the saturation concentration of water vapour, $D^0_{H_2O}$ and $D^0_{CO_2}$ are the standard diffusion coefficients of $H_2O$ and $CO_2$, and concentration difference of $CO_2$ inside and outside of the leaf is 0.0055 mol.m$^{-3}$ air, which is considered to be constant throughout the transition (Nobel, 2020; Töpfer et al., 2020).

In the first point, we have simulated the model using the constraints of $C_3$ metabolism (the ratio of carboxylase and oxygenase activity of RuBisCO is 3:1, there is no limitation on gaseous exchange etc.). In this step, the OP and aperture size throughout the three phases of day are assumed to be equal. Similar assumption is considered for the three phases of night as well. This refers to the 1st step of simulating the 1st point of the $C_3$-to-CAM continuum. However, in reality the stomatal opening and hence OP are not equal in different phases of day and night. To include this and to avoid the over-estimation of accumulation of osmolytes in any phase, we have recalculated the OP and aperture size depending on the $CO_2$ demand at each phase from the solution space. Using these newly calculated OP and aperture size in different phases, we again simulate the model and get our final solution. This is the 2nd step of simulating the 1st point of the $C_3$-to-CAM continuum. From this 1st point, keeping the new calculated aperture and OP fixed, we simulate the model at a reduced water loss. Again, we recalculate the OP and aperture size at each phase and simulate the model to get the final solution for the 2nd point of the $C_3$-to-CAM continuum. This two-step iterated simulation is repeated until the water loss becomes 600 µmol.m$^{-2}$.s$^{-1}$ and we reach to CAM.

In reality, water loss through stomata is dependent on the opening of the stomatal pore and hence the gaseous exchange occurring through the pore. Consequently, our calculations account for both $CO_2$ uptake and release, rather than focusing solely on $CO_2$ demand (as in Töpfer et al., 2020). For each phase, we calculate water loss based on whichever is greater: $CO_2$ uptake or $CO_2$ release (For simplicity we have included the exchange of $CO_2$ only, not of $O_2$). Additionally, we consider 300 MC and 1 GC in these calculations. As Reckmann et al., 1990 reported a basal level OP and aperture at nighttime, we have considered that the aperture is never completely closed and there is always a basal level opening of stomata. We have used minimization of total cellular flux as the objective function for each point throughout the transition and the phloem output (**Supplementary Table S1**) is kept constant throughout the transition.

From the solution space of each point in the $C_3$-to-CAM continuum (Provided in **Supplementary Data S2**), we calculate the Spearman's correlation (also called rank correlation) between the flux through each reaction and the water loss throughout the transition to get a list of reactions whose activities gradually increase or decrease along the $C_3$ to CAM transition. Ratio of carboxylase and oxygenase activity of RuBisCO is set to increase gradually from 3 to 5.15 (as in Tay et al., 2021). From the solution of the $C_3$ point, we find that the ratio of daytime $CO_2$ uptake and $O_2$ release is 1.23, which is experimentally reported to be nearly 1 (Canvin et al., 1980). This ratio is maintained along the $C_3$ to CAM transition to ensure that a decrease in pore size results in a proportional reduction in both $CO_2$ uptake and $O_2$ release. All the other constraints are given in **Supplementary Table S1.**

**Supplementary Table S1**. Constraints applied to GC and MC throughout the $C_3$-CAM transition

| Type of Cell | Constraints | Values |
|---|---|---|
| MC | Photon ratio in the three phases of day | 5:148:3 |
| | Nitrate uptake ratio at day and night* | 3:2 |
| | Phloem output rate (in $\mu mol.m^{-2}.s^{-1}$) | 0.259 |
| | Phloem output ratio at day and night** | 3:1 |
| | % of total accumulated osmolyte in GC to be sucrose, transferred from MC to GC in phase 2 at daytime | 80 |
| | % of total accumulated osmolyte in GC to be sucrose, transferred from MC to GC in phase 4 at nighttime | 80 |
| | C:O of RuBisCO in each phase of day in $C_3$ | 3:1 |
| | C:O of RuBisCO in each phase of day in CAM | 5.15:1 |
| GC | Photon ratio in the three phases of day | 5:148:3 |
| | Nitrate uptake | 0 |
| | Phloem output | 0 |
| | C:O of RuBisCO in each phase of day in $C_3$ | 3:1 |
| | C:O of RuBisCO in each phase of day in CAM | 5.15:1 |

*Further distributed according to the length of the phases

**Free to be produced in any phase of day and night

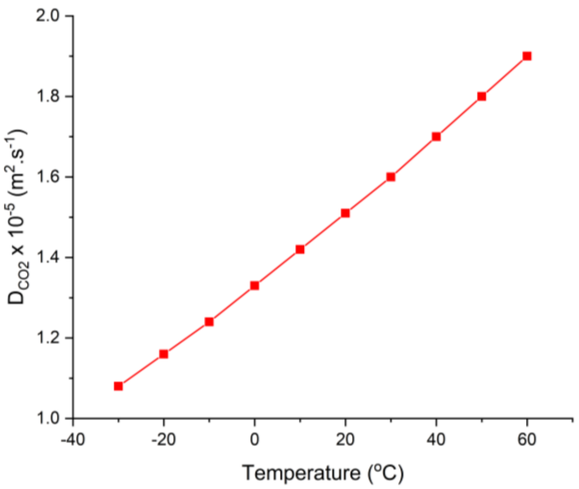


**Supplementary Figure S1**: Plot of the diffusion coefficient of $CO_2$ ($D_{CO_2}$) vs temperature (T)

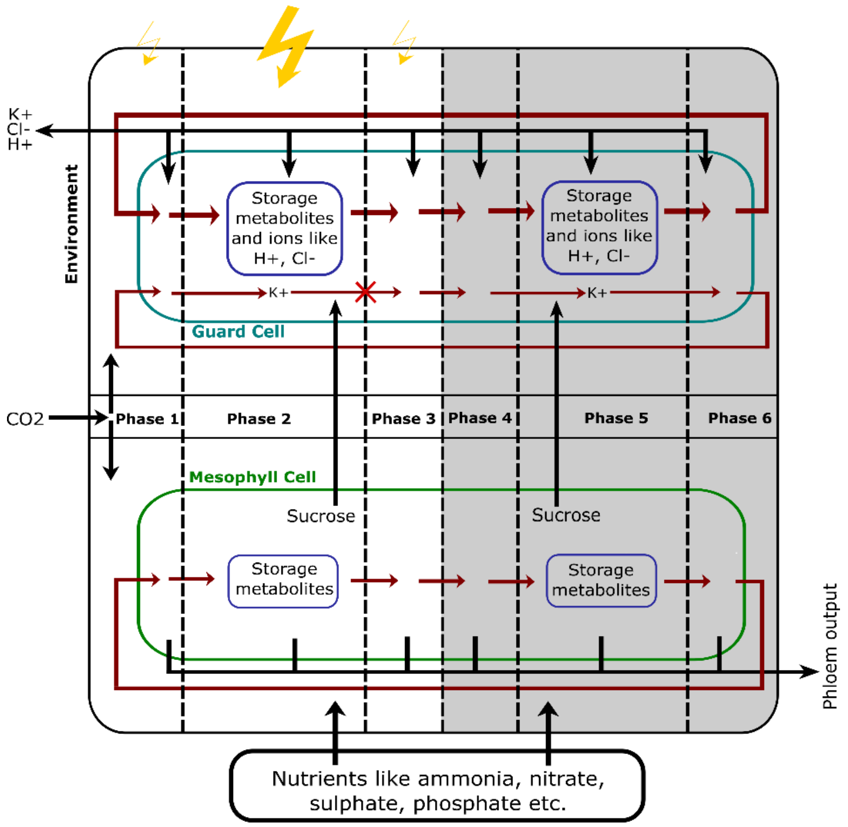


**Supplementary Figure S2**: Schematic representation of the model.

**Details related to Figure 1b**

Results show that as the transpirational water loss throughout the diel cycle decreases, the $C_3$ metabolism gradually shifts towards CAM. In $C_3$ metabolism, MC exhibits minimal starch and malate storage during the day. In contrast, CAM metabolism is characterized by substantial starch storage during the day and malate storage at night, as reported by Winter and Smith, (2022). GC shows a distinct metabolic pattern. In $C_3$, GC shows 3.6 times more starch storage during the day compared to MC, and starch production is also observed during the first phase of the night (phase 4). Whereas in CAM, daytime starch storage increases, but no starch synthesis is noted during the night. However, a small amount of malate storage is observed in CAM GC during the night. **Figure 1b** shows how the daytime starch and nighttime malate storage increased in GC and MC during the $C_3$ to CAM transition.

**Details related to Figure 1c**

In $C_3$, the stomata open at day and hence the OP is majorly maintained at day to balance the turgor pressure inside GC by accumulating the osmolytes. Now, as the metabolism shifts from $C_3$ to CAM, OP starts to decrease gradually at midday and increase at night depending on the time of $CO_2$ uptake (as described previously). However, at the dawn (phase 1) and afternoon (phase 3), the OP initially increases and then decreases as the metabolism shifted from weak to strong CAM. Kong et al., 2020 showed that stomatal aperture decreased at day and increased at night due to salt stress.

To balance the OP at day and night, the time of uptake and release of $K^+$ changes throughout the transition. In $C_3$, $K^+$ enters GC in phase 1 and leaves in phase 2, which represent the accumulation of $K^+$ at early morning (Lawson, 2009). Whereas, in CAM, $K^+$ starts entering the GC in phase 5, with its flux increasing in phase 6. This supports the fact that to take up maximum $CO_2$ just before dawn, a large OP and hence a large accumulation of osmolytes are necessary in the last phase of night. However, the maximum amount of $K^+$ comes out of GC in phase 1, which results in a limited amount of $K^+$ being stored in phase 1 and released in phase 2 to support a little $CO_2$ uptake at dawn in CAM. At the intermediate stage of transpirational water loss, $K^+$ enters the GC in both phase 6 and phase 1 and releases in phase 2. During the transition, the uptake of $CO_2$ continuously increases at midnight and last phases of night and decreases at midday. Whereas, at dawn it firstly increases and then decreases. To support this pattern of $CO_2$ uptake and aperture opening, the uptake of $K^+$ is increased in the mid night (phase 5) and the last phase of night (phase 6) to maintain higher osmolyte accumulation. Whereas in phase 1, we observe a sudden change in the pattern of $K^+$ uptake and release. At first the intake of $K^+$ in phase 1 increases with increase in OP and after a certain point, instead of up-taking the ions, $K^+$ transporter starts to release $K^+$. **Figure 1c** shows how the time of uptake and release of $K^+$ changes throughout the diel cycle along the transition. The subgraph in **Figure 1c** illustrates the temporal variations in $K^+$ uptake and release at the intermediate points between the final two data points in the main graph. This demonstrates that, at these last points, $K^+$ uptake and release occur change gradually but rapidly over time. The steeper appearance of the graphs in this short transition period reflects the high rate of change.

**Details related to Figure 1d-i and Figure 2**

Previous experimental studies have analysed up- and down-regulation of enzymes in $C_3$, CAM and their intermediate species during day and night (Heyduk et al., 2019), adaptation of CAM by plants due to draught-stress (Cushman and Borland, 2002), changes in enzyme activities at day and night in GC of ice plant due to the transition from $C_3$ to CAM by introducing salinity stress (Kong et al., 2020) and compared the proteomics of GC and MC during the transition from $C_3$ to CAM (Guan et al., 2021). However, the differentially expressed enzymes may not remain active continuously throughout the entire day or night. Our study encompasses six phases of the diel cycle, and the results not only corroborate previously observed patterns of enzyme expression during day and night but also reveals the activity of these enzymes at various phases throughout the diel cycle.

**Figure 1d-i** show that the enzyme activities in GC and MC can vary not only between day and night but also throughout different phases of the diel cycle. **Figure 2** shows the rank correlation values between water loss and the flux through each reaction throughout the $C_3$ to CAM transition in GC and MC for each phase. This helps us in identifying the enzymes and transporters which exhibit gradual changes throughout the $C_3$ to CAM transition and in which specific phases these changes occur.

1. The transition from $C_3$ to CAM results in higher daytime production and storage of starch and nighttime storage of malate in both GC and MC (**Figure 1b**). This increase in starch storage reflects the greater need of starch to supply PEP at night through glycolysis for dark $CO_2$ fixation, a well-known feature of CAM. This PEP is utilized in fixing nighttime $CO_2$ through PEPC into oxaloacetate which is further converted into malate by cytosolic MDH. This malate is stored at night and gets decarboxylated at daytime by PEPCK or NADP-ME in both the cells to provide $CO_2$ at daytime. We observe that in $C_3$, PEPC activity is higher in GC during the day (Phase 1) and night (Phase 4) compared to MC (**Figure 1d, g**) as reported in Daloso et al., 2017. As the metabolism shifts towards CAM, PEPC activity increases in midnight (phase 5) and at the end of night (phase 6) in GC (**Figure 1f, 2**), whereas throughout all phases of night in MC (**Figure 1i, 2**). Cytosolic MDH shows similar patterns in GC and MC. These results are in strong agreement with experimental findings- i) Guan et al. (2021) showed gradually increased protein abundance of cytosolic MDH, PEPC during the transition in both the cells with a greater increment in MC due to salt stress, ii) Cushman and Borland, (2002) reported the upregulation of the PEPC and MDH activity at night in MC by water limitation and, iii) Kong et al., (2020) showed PEPC activities at 2 am and 4 am in GC during the $C_3$ to CAM transition due to salt stress.

2. The stored malate can be decarboxylated during the daytime by two enzymes: NADP-ME and PEPCK. A correlation study shows that PEPCK activity during the transition increases in phase 2 for both GC and MC (**Figure 1f,i and Figure 2**), while no correlation is observed for ME. However, when examining the correlation of the combined activity of ME and PEPCK with water loss in phase 2, the correlation is 1 (**Figure 1d-i)**. This indicates that, during the transition

from $C_3$ to CAM, both ME and PEPCK work together to decarboxylate malate. There is an increase in protein abundance of NADP-ME, observed in Guan et al., (2021), which is in accordance with our results.

3. To maintain the OP at daytime in GC, MC supplies sucrose to GC during phase 2, accounting for 80% of the total osmolytes required for maintaining OP (described earlier). The remaining OP is maintained by sucrose production within the GC itself. **Figure 2** shows that the activity of sucrose synthase is downregulated in GC and MC during phase 2 as the metabolism shifts towards CAM. This reduction in sucrose synthase activity explains the decreased requirement for sucrose production in both MC and GC at daytime as the metabolism shifts towards CAM.

4. Pentose phosphate pathway (PPP) shows differential activities across different phases in GC and MC. Guan et al. (2021) showed that the protein abundance of transketolase, an enzyme of oxidative PPP (OPPP), is increased in GC and decreased in MC. Tay et al., 2021 also reported the decreased flux of OPPP at night in MC during the $C_3$ to CAM transition. Our results reveal that the three OPPP enzymes- transketolase 1, transketolase 2, and transaldolase—are downregulated in MC throughout all three phases of night. In contrast, GC shows increased enzyme activity in phases 2 and 4 but decreased activity in phase 1. Nonetheless, the total activities across the day of these enzymes are increased in GC. However, E4P takes an alternative path using SBPaldolase and SBPase to get converted to S7P in phase 2 in GC and phase 3 in MC (Sharkey, 2021). Three enzymes of non-oxidative part of PPP (G6PDH, 6PGD and 6PGL) also show decreased activity throughout all the phases of night and the 1$^{st}$ phase of day in MC.

5. GC and MC both produce starch through gluconeogenesis and degrade starch through glycolysis. Guan et al. (2021) have reported changes in protein abundance of different enzymes of glycolysis, gluconeogenesis, ETC, TCA cycle, Calvin cycle, PPP etc., in GC and MC due to their metabolic shift from $C_3$ to CAM. But the activities of these enzymes are expected to change differently in these cells in different phases throughout the diel cycle. The observed increased activities of the enzyme related to gluconeogenesis such as Enolase, PGM, PGK, G3PDH, FBPA, FBPase, PGI etc in chloroplast and cytosol in phase 2 in GC and MC support the increased starch production in CAM. Increased activities of glycogen synthase and other enzymes related to starch production are also observed in phase 2 in both the cells. Guan et al., 2021 showed an increase in the abundances of enolase, G3PDH and FBPA, which matches our result. Whereas the starch breakdown through glycogen phosphorylase is observed to be increased in midnight and last phase of night in GC and throughout all phases of night in MC. Kong et al., (2020) showed increased expression of *GTF1*, a gene involved in starch

degradation, at 2 am and 4 am, which supports our results. Glycolytic enzymes like PGI, PFK, FBPA, TPI, G3PDH, PGK, PGM, Enolase, PK etc., show increased activity in all the phases of night in MC and phase 5 and 6 in GC. This supports- i) the necessity of conversion of starch into PEP for providing substrate to PEPC and ii) the increased supply of carbon skeleton to TCA cycle to increase the ATP production through ETC (Tan and Cheung, 2020). However, at dusk (phase 4) metabolism changes in a different manner in GC. In $C_3$ GC, besides daytime, starch synthesis occurs at the dusk (phase 4) also. Sucrose, which is the major osmolyte in balancing OP at mid-day is used in producing starch via gluconeogenesis and the dark $CO_2$ fixation occurs through PEPC in phase 4. As soon as the metabolism starts shifting from $C_3$, GC loses this property. This infers that as we shift towards CAM, the $CO_2$ fixation starts happening majorly at mid-night and the refixation of this $CO_2$ into starch through gluconeogenesis occurs at daytime. Carbonic anhydrase is also upregulated in all phases of night in MC and in phase 5 and 6 in GC.

6. Moreover, the enzymes of TCA cycle show differential activities in GC and MC. PDH and CS show increased activity at midnight and the end of night in GC and throughout all phases in MC. Rest of the enzymes majorly show increased activity in phase 2, 5 and 6 in MC and phase 2 in GC and decreased activity in phase 4 and 5 in GC. Complexes of ETC- NADH dehydrogenase (Complex I), cytochrome $bc_1$ (Complex III), cytochrome c oxidase (complex IV) and ATP synthase (Complex V) also show increased activities in all the 6 phases in MC and phase 2 and 6 in GC, which justifies the increase in ATP demand both at day and night as the metabolism shifts towards CAM. Guan et al., 2021 also reported an increase in the abundances of citrate synthase, NADH dehydrogenase (complex I) and cytochrome c oxidase (complex IV) in both GC and MC. Moreover, in GC, nighttime citrate storage plays an important role in providing reductant through TCA cycle for ATP production through ETC at daytime. The stored citrate is utilized in TCA for the production

7. During this transition, the activities of different mitochondrial-cytosolic and plastidial-cytosolic transporters adjust to support shifts in central carbon metabolism. These include GAP and G6P transporters and OAA-Mal, GLT-Mal and 2KG-Mal shuttles in plastid and OAA-Mal and 2KG-Mal shuttles in mitochondria. During the day, GAP produced in the GC is used in the oxidative pentose phosphate pathway (OPPP). Meanwhile, GAP produced in the MC during the day moves to the cytosol, where it is converted to F6P via gluconeogenesis enzymes and then transported to the plastid for starch production. Consequently, as starch production in the MC increases, the flux through these transporters also rises. Similarly, at night, as starch breakdown intensifies in both cells. The flux through the plastidial-cytosolic G6P transporter increases to facilitate the movement of G6P to the cytosol for glycolysis. The plastidial-

cytosolic Glt-Mal and 2KG-Mal shuttles facilitate the transport of 2KG into the plastid, where it is used to synthesize Glt. This Glt is then transported to the cytosol. Although, Guan et al., 2021 reported decreased in abundances of glutamate synthase in GC and MC, our results show increased activity of in GC and decreased in MC.

The Mal-OAA shuttle supplies OAA to the plastid and generates NADP by converting OAA to Mal through plastidial MDH. Consequently, the flux patterns of MDH and the Mal-OAA shuttle are similar. On the other hand, the mitochondrial-cytosolic Mal-OAA and Mal-2KG transporters deliver malate produced in the cytosol during both day and night, thereby enhancing the production of reductants (NADH) needed for increased ATP synthesis. Therefore, we can conclude that the shuttles work in different manners for balancing redox in mitochondria, chloroplast and cytosol.

In summary, enzymatic activities of central carbon metabolism and the flux through transporters vary across different phases of the day and night. Phase 4 shows distinct changes in enzyme activity compared to phases 5 and 6, while major alterations during daytime are observed in phases 1 and 2. This study indicates that GC has a unique pattern of metabolic transition to CAM. Therefore, incorporating engineering strategies to modify GC metabolism, in addition to MC, could assist biotechnologists in introducing CAM characteristics into $C_3$ plants.

**Result using different temperature and relative humidity**

| | $T_{Max}$ (°C) | $T_{Min}$ (°C) | $RH_{Max}$ | $RH_{Min}$ | Total water loss for $C_3$ ($\mu$mol.m$^{-2}$.s$^{-1}$) | Water loss for CAM ($\mu$mol.m$^{-2}$.s$^{-1}$) |
|---|---|---|---|---|---|---|
| Töpfer et al., 2020 | 30.0 | 15.0 | 1.000 | 0.400 | 13,42,711 | 1,80,280 |
| Jaipur, India | 44.6 | 24.9 | 0.961 | 0.133 | 40,85,944 | 4,29,798 |

The chart illustrates how the amount of transpirational water loss varies in $C_3$ and CAM plants based on the environmental factors like temperature (T) and relative humidity (RH). The maximum and minimum values of temperature ($T_{Max}$ and $T_{Min}$) and relative humidity ($RH_{Max}$ and $RH_{Min}$) of Jaipur are based on the measurements of IMD in Jaipur, India. Given that Jaipur's climate is extremely hot and dry, $C_3$ plants experience significantly higher water loss. Even at the extreme end of the $C_3$-to-CAM continuum, water loss remains much higher compared to the water loss we get using the values of T and RH of Germany provided in Töpfer et al., 2020. While total water loss for CAM in Germany is nearly 1/8$^{th}$ of that of $C_3$, the value is 1/10$^{th}$ in case of Jaipur. Additionally, previously we observed the dark time fixation of $CO_2$ by PEPC in GC in phase 4. Whereas, in case of Jaipur, this activity is lost and the starch production relies on stored sucrose (Tan and Cheung, 2020). While the overall pattern of metabolic changes during the transition is nearly identical in both cases, there are some subtle differences that suggest the activation of different alternative pathways to achieve the goal.
